# Generative Diffusion Models for Antibody Design, Docking, and Optimization

**DOI:** 10.1101/2023.09.25.559190

**Authors:** Zhangzhi Peng, Chenchen Han, Xiaohan Wang, Dapeng Li, Fajie Yuan

**Affiliations:** Electrical Engineering & Computer Science, University of Missouri, Columbia, 65211, MO, USA; School of Life Engineering, Westlake University, Hangzhou, 310024, Zhejiang, China; Center for Infectious Disease Research, Westlake University, Hangzhou, 310024, Zhejiang, China; School of Life Sciences, Westlake University, Hangzhou, 310024, Zhejiang, China

## Abstract

In recent years, optimizing antibody binding affinity for biomedical applications has become increasingly important. However, traditional wet-experiment-based approaches are time-consuming and inefficient. To address this issue, we propose a diffusion model-based antibody optimization pipeline to improve binding affinity. Our approach involves two key models: AbDesign for designing antibody sequences and structures, and AbDock, a paratope-epitope docking model, used for screening designed CDRs. On an independent test set, our AbDesign demonstrates the exceptional performance of an RMSD of 2.56Å in structure design and an amino acid recovery of 36.47% in sequence design. In a paratope-epitope docking test set, our AbDock achieves a state-of-the-art performance of DockQ 0.44, irms 2.71Å, fnat 0.40, and Lrms 6.29Å. The effectiveness of the optimization pipeline is further experimentally validated by optimizing a flaviviruse antibody 1G5.3, resulting in a broad-spectrum antibody that demonstrates improved binding to 6 out of the nine tested flaviviruses. This research offers a general-purpose methodology to enhance antibody functionality without training on data from specific antigens.

## 1 Introduction

Antibodies play crucial roles in the immune system, exhibiting exceptional specificity and affinity for specific antigens. Therefore, the optimization of antibody binding affinity is of paramount importance in a wide range of biomedical applications, including therapeutics, diagnostics, and research. The binding affinity is primarily determined by the complementary-determining regions (CDRs), among which the third CDR of the heavy chain (CDRH3) contributes significantly to the antibody-antigen interaction.

The conventional step-wise approaches in antibody design and optimization, while yielding valuable insights, can be exceedingly time-consuming, and considerable effort may be expended on interrogating antibodies that ultimately prove nonfunctional [1–3]. Furthermore, when improved binding is achieved, there may arise the need for subsequent modifications to enhance other properties, such as hydrophobicity [4]. These subsequent alterations can potentially have a detrimental impact on the previously optimized binding, necessitating additional measurement and engineering cycles. In practice, the process of identifying the final antibody candidate routinely consumes more than a year. This extended timeline is mainly due to the intricate and labor-intensive nature of these iterative procedures [5].

In recent years, machine learning (ML) models have demonstrated remarkable potential in the development of antibodies with enhanced binding characteristics. [6–9]. Recent works [10–12] utilize geometric graph neural networks trained on antibody-antigen complex data to predict the CDR sequences and structures. Despite their impressive in-silico results, the designed CDRs are not experimentally verified. Notably, a study by [13] introduced a novel deep learning framework that successfully broadens the neutralizing activity against diverse SARS-CoV-2 variants by learning the mutational effect on protein-protein interactions from protein complex structures and binding affinity data. However, these methods rely on obtaining binding affinity data for specific targets—a process that is often prohibitively time-consuming, labor-intensive, and costly. Addressing this challenge, recent research by [14] utilizes language models trained on extensive protein sequence datasets to propose mutations within antibody sequences. This method offers a general approach for enhancing the binding affinity of antibodies across multiple targets, relying solely on antibody sequences as input. However, this approach does not take into account the structural interactions occurring at antibody-antigen interfaces. Consequently, there is a potential risk of reduced specificity. As reported in their study, approximately half of the substitutions recommended by the language model (including around half of the substitutions) are located within framework regions that may not contribute to the desired binding.

In this study, we introduce an antibody optimization approach that combines two stages, antibody design and screening, involving AbDesign and AbDock. These two models leverage the generative capabilities in diffusion models, which have demonstrated impressive performance in generating images [15–18], language [19], and biomolecules [10, 20]. During the design phase, AbDesign takes advantage of its generative nature to explore the vast sequence space, which is crucial for the identification of optimal antibody sequences. In the subsequent screening stage, we employ a paratope-epitope docking model, which we trained to generate paratope docking poses to a given epitope. While our approach does not explicitly train a scoring model for binding affinity prediction, AbDock can assess the quality of antibody candidates by evaluating the generated docking poses, which serve as a proxy for antibody scoring. Both AbDesign and AbDock have been trained using antibody-antigen complex structures available in the PDB database. This approach eliminates the need for labor-intensive wet experiments to generate training data and enables our method to generalize effectively across diverse antigens.

To showcase the efficacy of our AbDesign approach in antibody design, we conducted a comprehensive evaluation using a meticulously curated test dataset. In this evaluation, AbDesign demonstrated remarkable performance, attaining an RMSD (Root Mean Square Deviation) of 2.56Å in structural design and an impressive amino acid recovery of 36.47% in sequence design. In a separate set of tests for paratope-epitope docking, our AbDock method showcased state-of-the-art capabilities, achieving outstanding results across various evaluation metrics. Notably, it achieved a DockQ score of 0.44, an irms (interface root mean square) of 2.71Å, a fnat (fraction of native contacts) score of 0.40, and an Lrms (ligand root mean square) of 6.29Å. These results underscore the exceptional performance of our AbDock approach in accurately predicting antibody-antigen interactions. To further exemplify the efficacy of our methodology, we undertook the optimization of the monoclonal antibody 1G5.3. This particular antibody, denoted as 1G5.3, has demonstrated a notable affinity and crossreactivity towards the NS1 proteins that are synthesized by multiple flaviviruses [21]. Consequently, the monoclonal antibody 1G5.3 holds the promise of conferring a comprehensive shield against a spectrum of flavivirus infections. By designing four residues in the CDRH3 of 1G5.3, we successfully obtained and experimentally validated a variant that exhibited an exceptional and expansive binding capacity, effectively targeting six out of the nine tested flaviviruses. This remarkable outcome attests to the variant’s potential as an immunotherapeutic agent for offering universal protection against flavivirus infections. By capitalizing on the capabilities of generative diffusion models and integrating AbDesign and AbDock, our approach offers a general-purpose framework for optimizing antibody binding affinity without affinity data for specific targets.

## 2 Results

In this study, we introduce two novel models, namely AbDesign and AbDock, designed for antibody CDR sequence-structure co-design (Fig. 1 A) and paratope-epitope docking (Fig. 1 B), respectively. The co-design task focuses on predicting the CDR sequence and conformation given the known paratope-epitope binding interface. Conversely, the docking task aims to predict the binding conformation of the paratope when the epitope pocket and paratope sequence are known. Refer to Section 4.1 for a more rigorous task definition. Both AbDesign and AbDock models are built upon the foundations of diffusion models. We follow the formulation of existing diffusion models [10, 22] and define the generation of CDR as a denoising process applied to protein backbone frames (*C*_*α*_ position and backbone orientation), as shown in Fig. 1 C. To accomplish this, we train a neural network capable of effectively denoising the data. We recognize that the two tasks involved possess distinct characteristics, which necessitate the use of different neural network architectures. For the CDR sequence structure co-design task, where the binding interface is provided and only the CDR region remains unknown, we propose the adoption of the Multi-Channel Equivariant Neural Network (MC-EGNN) [23–25], shown in Fig. 1 D left. This network effectively models the contextual information of the binding interface as a locally-connected graph, facilitating the accurate prediction of the CDR while remains low computation and memory efficiency. In the case of the paratope-epitope docking task, where solely the epitope pocket structure is given, we employ a more expressive neural network called the invariant point attention (IPA) network [26], shown in Fig. 1 D right. This network is adept at capturing the intricate structural complexities of the system, modeling it as a fully-connected graph. In Fig. 1 E, we present the antibody optimization pipelines by using the proposed AbDesign and AbDock, where the AbDock is used to generate binding poses and in-silico scoring, while the AbDesign is employed for sequence design.

**Fig. 1.**
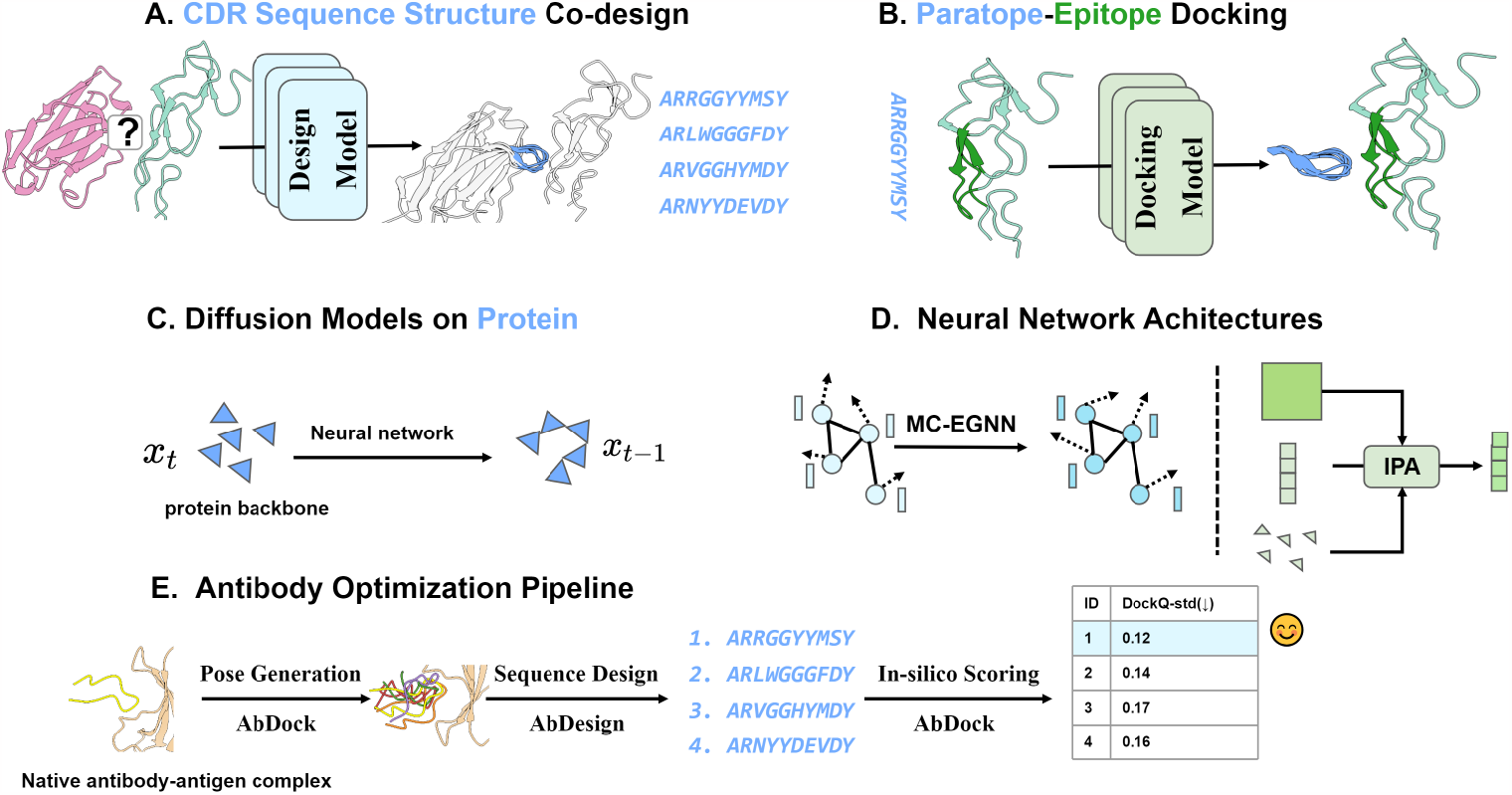
Overview of proposed methods. A: Flowchart of the CDR sequence structure co-design task. B: Flowchart of the paratope-epitope docking task. C: The formulation of diffusion model defined on protein backbones. D: The illustration of two neural networks: Multi-Channel Equivariant Neural Network (MC-EGNN) (left) and Invariant Point Attention (IPA) (right). E: The flowchart of the proposed antibody optimization pipeline.

In order to validate the effectiveness of AbDesign and AbDock, a comprehensive performance evaluation was conducted on separate held-out design and docking test sets, as outlined in Section 2.1 and 2.2. The results obtained from these evaluations demonstrate that our proposed AbDesign and AbDock models surpass state-of-the-art baselines across several metrics, indicating their superior performance in the respective tasks.

In Section 2.3, we undertake a detailed evaluation of the model’s prediction capacity, revealing the model’s strength and weakness and highlighting its robustness. In addition, we compare the designed structures and sequences with native counterparts, revealing that the sampled sequences and structures possess a natural resemblance to native CDRs.

One promising application of paratope-epitope docking is virtual screening, which enables the efficient and cost-effective screening of large-scale potential antibody candidates against a specific target. In Section 2.4, we provide empirical evidence where the scoring results generated by AbDock exhibit a discernible correlation with experimentally-determined affinities. This correlation underscores the viability of our AbDock as a general screening tool for computational antibody design.

Lastly, in section 2.5, we apply the antibody optimization pipeline to the monoclonal antibody 1G5.3 and show the optimized antibody exhibiting improved binding affinity on six out of the nine tested flaviviruses.

### 2.1 Performance Evaluation of AbDesign on Design Task

We assess the effectiveness of our AbDesign on the CDR design tasks by comparing performance with state-of-the-art baselines. The evaluation aims to validate the model’s ability to accurately predict CDR sequences and conformations when the paratope-epitope binding interface is known. We compared with existing CDR design methods as follows. DiffAb [10], DiffAb is a diffusion model-based antibody sequence-structure co-design method. AR-GNN [11], AR-GNN is an autoregressive graph generation model, which generates an antibody graph residue by residue. MEAN [12], MEAN is a graph-based model for CDR sequence-structure co-design. RefineGNN [11], RefineGNN is an iterative refinement graph neural network that co-designs the sequence and 3D structure of complementarity-determining regions (CDRs) as graphs. We additionally trained a fixed backbone sequence design version of AbDeisn, named AbDesign (seq. design), where the CDR structure is given, and the model only predicts the sequence. Following [10], we collect antibody-antigen data from the SAbDab database [27]. We first remove structures whose resolution is worse than 4Å and remove antibodies targeting non-protein antigens. We cluster antibodies in the database according to CDR-H3 sequences at 50% sequence identity using MMseqs2 [28]. we selected five independent clusters representing diverse antigen types. These clusters comprised 19 complex structures featuring antigens from well-known pathogens such as SARS-CoV-2, MERS, influenza, and others. The remaining structures from the dataset were utilized for model training. This evaluation encompassed the following metrics. 1) Root Mean Square Deviation (RMSD): RMSD calculates the average deviation between the designed CDR structures and the corresponding native structures. 2) Amino Acid Recovery (AAR), is the percentage of designed amino acids identical to native amino acids.

In Figure 2 A, we present the performance comparison of structural design, wherein our AbDesign achieves an RMSD of 2.56Å, surpassing AR-GNN and DiffAb by over 1Å, and even outperforming RefineGNN by 0.13Å. Notably, AbDesign also exhibits greater stability across various test cases, as indicated by its lower variance compared to the baseline models. Moving to Figure 2 B, we display the comparison of sequence design. Here, our AbDesign (seq. design) achieves 36.47%, while the overall AbDesign performance reaches an impressive 49.55%. These values significantly surpass the results obtained from all baseline methods, underscoring the substantial advantage of our approach. In Figure 2 C, we visually present the designed CDR sequence and structure of the BD-604 Fab (PDB code: 7chf), a key binder to the SARS-CoV-2 RBD. Evidently, our AbDesign methodology yields a near-native conformation with an RMSD of 1.22Å, accompanied by a noteworthy amino acid recovery of 63%. In contrast, it’s worth noting that DiffAb designs a CDR characterized by a beta-sheet structure, while AR-GNN and RefineGNN generate CDRs containing bond breaks.

**Fig. 2.**
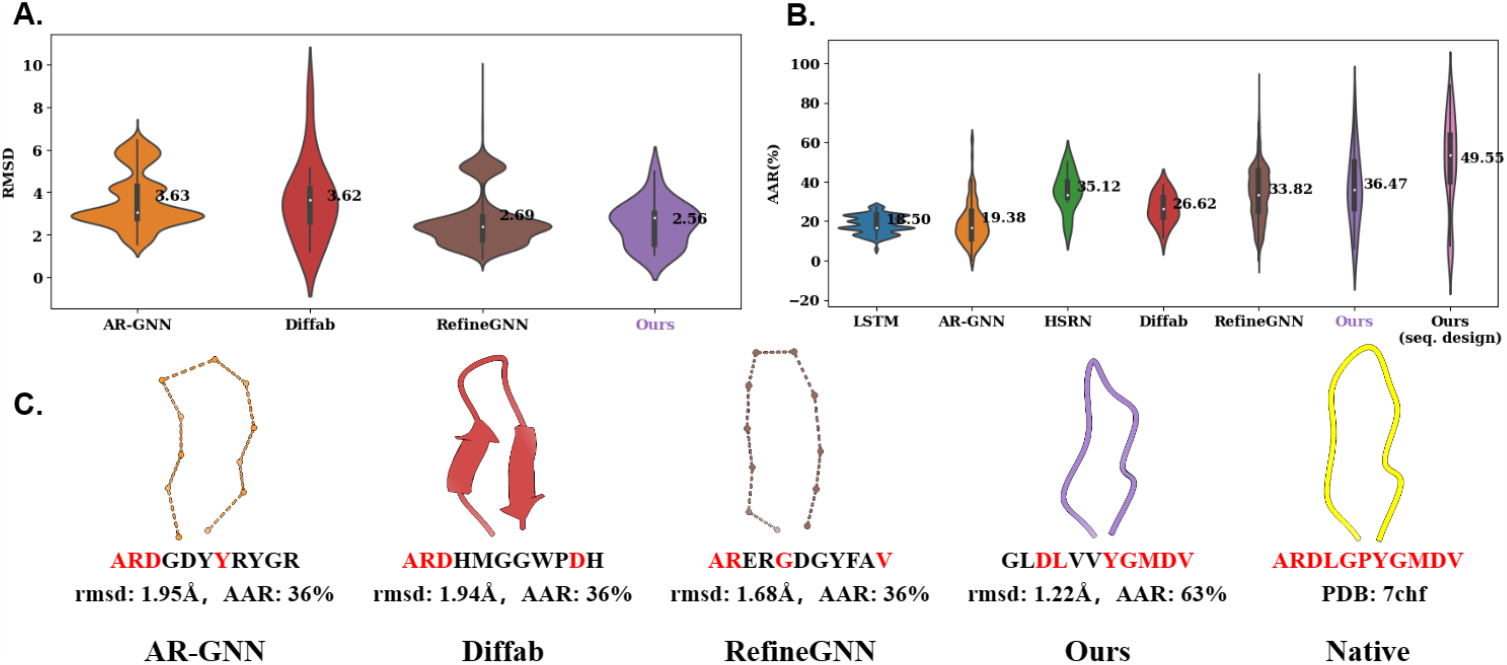
Performance comparison on the antibody design task. This figure presents a comparative analysis of CDR design performance, including both structural design performance (A) evaluated using Root Mean Square Deviation (RMSD), sequence design performance (B) assessed by Amino Acid Recovery (AAR), and a visualization (C) showcasing the CDR designs generated by different methods.

### 2.2 Performance Evaluation of AbDock on Docking Task

To evaluate the model’s ability to accurately predict the binding pose of the paratope given the sequence and epitope pocket, we conducted a comprehensive evaluation on a separate held-out docking test set. We compare the AbDock with the following baselines. ZDOCK [29], ZDOCK is a protein–protein docking program based on the Fast Fourier Transform algorithm. HDOCK [30], HDOCK is a hybrid algorithm of template-based and template-free protein–protein docking method. HSEN [31], HSRN is a graph-neural-network-based Antibody-Antigen Docking and Design method; we use the Docking model in HSRN for comparison. We also compare with the relaxed version of HSRN, HSRN (relaxed), where the predicted poses are relaxed using openMM [32]. We additionally train two versions of AbDock for exploration: AbDock (CDRs) for docking the CDRH1, CDRH2, and CDRH3, and AbDock (antibody) for docking the heavy chain. We select 44 samples from the processed data as the docking test set, and the rest non-overlapping data is used for training AbDock. We use DockQ as the evaluation metric to assess the accuracy of the docking pose prediction. DockQ is calculated by comparing the predicted docking poses with the native poses of the antibody-antigen complexes. It combines three distinct though related measures, namely Fnat: the fraction of native interfacial contacts preserved in the interface of the predicted paratope-epitope interface, LRMS, the Ligand Root Mean Square deviation calculated for the backbone of the paratope of the model after superposition of the epitope, and iRMS, the RMSD between predicted and native interface residues. The three metrics are also reported. For each test case, we generate 100 docking poses using AbDock and select one for evaluation. We cluster the 100 generated poses in terms of structural similarity (evaluated by RMSD) and select the center pose.

In Fig. 3, we present a performance comparison of DockQ, irms, fnat, and RMS using a violin plot (Fig. 3, A B C D). Our model, AbDock (HCDR3), achieves the highest DockQ score of 0.44, outperforming the second-best model, HSRN, by 0.01.

**Fig. 3.**
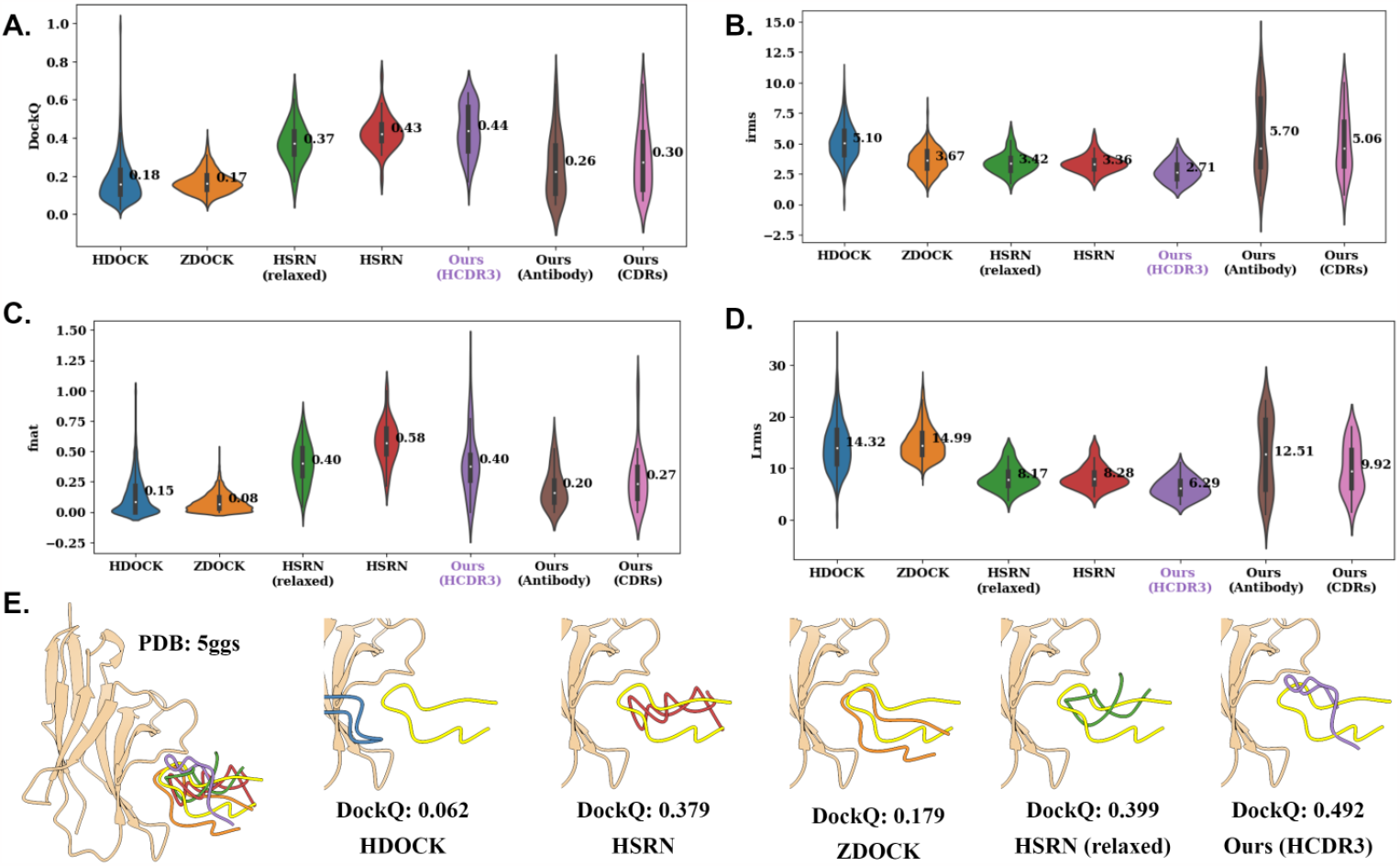
Performance comparison on paratope-epitope docking task. This figure presents a comparative evaluation of the docking performance for paratope-epitope interactions. Panels A, B, C, and D display the evaluation of DockQ, irms, fnat, and Lrms scores, respectively. Panel E showcases the visualization of docking poses generated by different methods.

Regarding irms, our model achieves a prediction accuracy of 2.71Å, surpassing the second-best model, HSRN (relaxed), by 0.71Å. This demonstrates our model’s superior capability in predicting interface structures. For fnat, our model achieves a score of 0.40, slightly lower than HSRN’s score of 0.58. Furthermore, in Lrms, our model achieves an error of 6.29Å, which is lower than the second-best model, HSRN (relaxed), by 1.88Å. However, comparing the distribution of Lrms, our AbDock consistently performs well across the test cases (centralized Lrms distribution). A comparative analysis between the performances of AbDock in different scenarios—specifically, AbDock (CDRs), AbDock (antibody), and AbDock (HCDR3)—reveals significant disparities. In the first two cases, the DockQ score is approximately half of that observed in the third case. This disparity underscores the increased difficulty inherent in docking the entire antibody-antigen complex as opposed to focusing solely on certain regions or components. In Fig. 3 E, we present the visualization of the predicted docking poses for the case PD-1 in complex with pembrolizumab Fab (PDB code: 5ggs), which is crucial in cancer immunotherapy. Our model generates the most native docking poses, with a DockQ score of 0.492, outperforming other methods, where their performance is relatively inferior. Notably, HDock fails to generate poses in the right pocket. Overall, our proposed AbDock (HCDR3) model demonstrates promising performance and outperforms existing methods in several critical metrics, making it a valuable candidate for accurate docking predictions.

### 2.3 Understanding Model’s Performance

#### Comparison of Model Performance at Fixed and Flexible Binding Pockets

In Fig. 4 A, we conducted a comprehensive comparison of our model’s performance in two distinct binding pocket scenarios: fixed and flexible pockets. The fixed pocket is characterized by the epitope forming secondary structures such as alpha-helices or beta-sheets, while the flexible pocket consists of loops. To assess the model’s performance for structure design, we employed the Root Mean Square Deviation (RMSD) metric, and for sequence design, we utilized the Amino Acid Recovery (AAR) metric. The results unequivocally demonstrate that our model excelled when the binding pocket was fixed. Specifically, in fixed pocket scenarios, the model exhibited a remarkable ability to learn and adopt the conformation of Complementarity-Determining Regions (CDRs), effectively integrating them into the pocket. This compelling finding indicates that the fixed pocket environment provided more favorable conditions for the model to achieve precise structure and sequence designs.

**Fig. 4.**
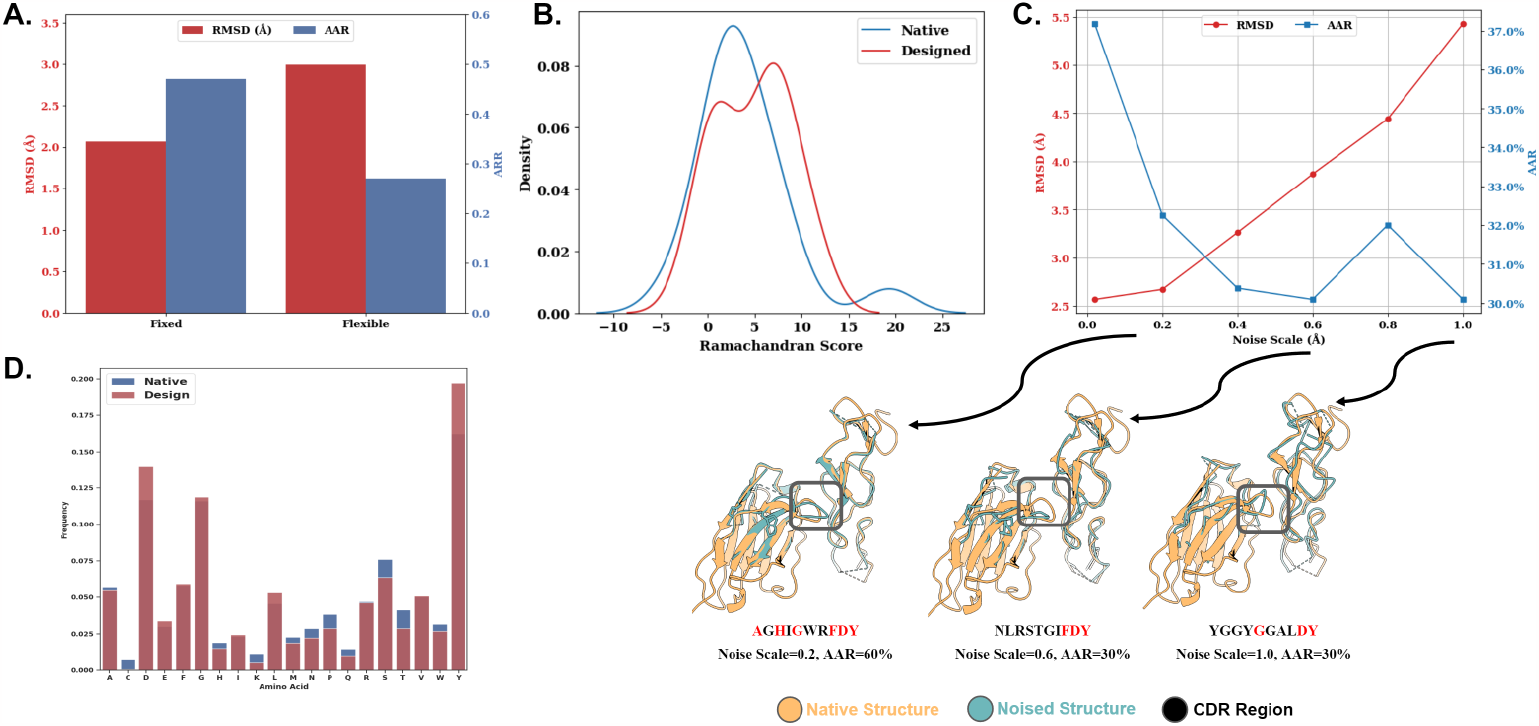
Performance analysis. This figure presents a comprehensive performance analysis. Panel A compares the model performance in fixed and flexible binding pockets. Panel B showcases the distribution of Ramachandran scores, comparing the designed CDR structures with the native ones. Panel C examines the robustness of sequence and structure design to input noise. Panel D presents a comparison of amino acid frequencies between native and designed sequences.

#### Ramachandran Score Distributions

Further analysis involved a comparison of the distribution of Ramachandran scores between the designed structures and native structures (Fig. 4 B). The Ramachandran score describes the distribution of (Φ, Ψ) torsion angles of the protein backbone and is a crucial indicator of the protein’s conformational quality. Remarkably, the comparison revealed a high degree of similarity between the distributions, suggesting that the designed structures closely resembled their native counterparts. This compelling result strongly supports the notion that our model was successful in generating antibody structures that align with the preferred conformational space found in native antibodies.

#### Robustness to Input Noise

To gauge the robustness of our model, we systematically introduced varying levels of noise to the input structure. By adding Gaussian noise at different scales to the input coordinates, we evaluated the sequence and structure design performance under different noise conditions. As shown in Fig. 4 C, the model’s performance showed a decline with increasing noise levels. However, it is noteworthy that even under high noise conditions, the AAR metric remained above 30%. This result indicates that the model could still generate relatively reliable sequence designs, showcasing a degree of resilience in the face of input noise. Nevertheless, it is important to acknowledge that the noise in the input data did have an adverse impact on the overall performance of the model.

#### Amino Acid Frequency Comparison

Lastly, we investigated the frequency distribution of amino acids in both the native and designed sequences (Fig. 4 D). Strikingly, the distributions demonstrated a high level of similarity, indicating that our model effectively captured the native-like amino acid preferences present in natural antibodies.

In conclusion, our experimental results underscore the remarkable performance of our model in fixed-binding pocket scenarios. Furthermore, the model demonstrated a certain level of robustness to input noise, with the AAR metric remaining relatively high under noisy conditions. Additionally, the similarity observed in the Ramachandran score distributions and amino acid frequencies further validate the native-like characteristics of the designed structures and sequences. These findings hold significant promise for the rational design of antibodies with desirable properties, offering valuable insights for advancing antibody engineering and therapeutic development.

### 2.4 Antibody Candidates Virtual Screening using AbDock

Virtual screening has emerged as a highly promising and cost-effective approach for efficiently screening a vast number of potential antibody candidates against a specific target. In this section, we present the application of paratope-epitope docking using AbDock as a practical and effective screening strategy for computational antibody design. While AbDock is primarily designed for docking and cannot directly score CDR sequence candidates, we can utilize the output docking metrics, such as DockQ, as a pseudo score. This approach is based on the assumption that a well-trained docking model has learned the intricate paratope-epitope interactions from native sequences. Consequently, poorly designed CDR sequences would result in inaccurate binding predictions, reflected by lower DockQ values.

To evaluate the performance of AbDock in virtual screening, we leverage the data from the study [33]. In this study, the authors assessed the binding affinities of 422 designed antibody CDRH3 variants towards the human epidermal growth factor receptor 2 (HER2). The dataset containing the designed CDRH3 sequences and affinity values, denoted as *− log*(*KD*(*M*)) (larger value denotes higher affinity), is publicly available at GitHub^1^. The native antibody-antigen complex structure of HER2 is available in Protein Data Bank (PDB) database with the PDB code 1N8Z^2^. By considering the designed CDRH3 sequences and the epitope pocket obtained from 1N8Z, we use AbDock to generate various docking poses. Subsequently, we computed the average (DockQ_avg) and standard deviation (DockQ_std) of the DockQ score for each designed CDRH3 sequence. The scores, namely DockQ_avg and DockQ_std, serve as indicators of the binding quality for a given CDRH3 sequence. A higher value of DockQ_avg indicates a higher likelihood of a designed CDRH3 sequence forming a favorable antibody-antigen interaction. Conversely, DockQ_std reflects the uncertainty associated with the AbDock prediction. Lower values of DockQ_std indicate a more confident prediction and correspond to CDRH3 sequences that are more likely to exhibit strong binding to antigens.

Fig. 5 (left) displays a scatter plot of these two variables, where we observed a positive correlation between DockQ_avg and *− log*(*KD*(*M*)) with Pearson’s correlation coefficient of 0.23 and Spearman’s correlation coefficient of 0.23. Regression result indicates that higher DockQ_avg suggests higher binding affinity with moderate correlation. Similarly, we analyzed the correlation between DockQ_std and *− log*(*KD*(*M*)) as shown in Fig. 5 (right). we found a negative correlation between DockQ_std and *− log*(*KD*(*M*)), with Pearson’s correlation coefficient of -0.29 and Spearman’s correlation coefficient of -0.27. The results verify that a well-trained model can not accurately dock a poor sequence and thus resulting in high standard deviation in DockQ (DockQ_std). Furthermore, the Figure presents four highlighted cases, each representing distinct binding affinity scenarios. the upper two cases with high binding affinity with high DockQ_avg and low DockQ_std, respectively, exhibiting favorable docking and high confidence in predictions. The docked poses in these cases exhibit consistent and stable orientations. Conversely, cases with low binding affinity display lower DockQ_avg scores and higher DockQ_std scores, displaying substantial structural variations and limited consensus in the predicted poses. Our model effectively distinguishes high-affinity sequences from low-affinity ones based on its prediction confidence.

**Fig. 5.**
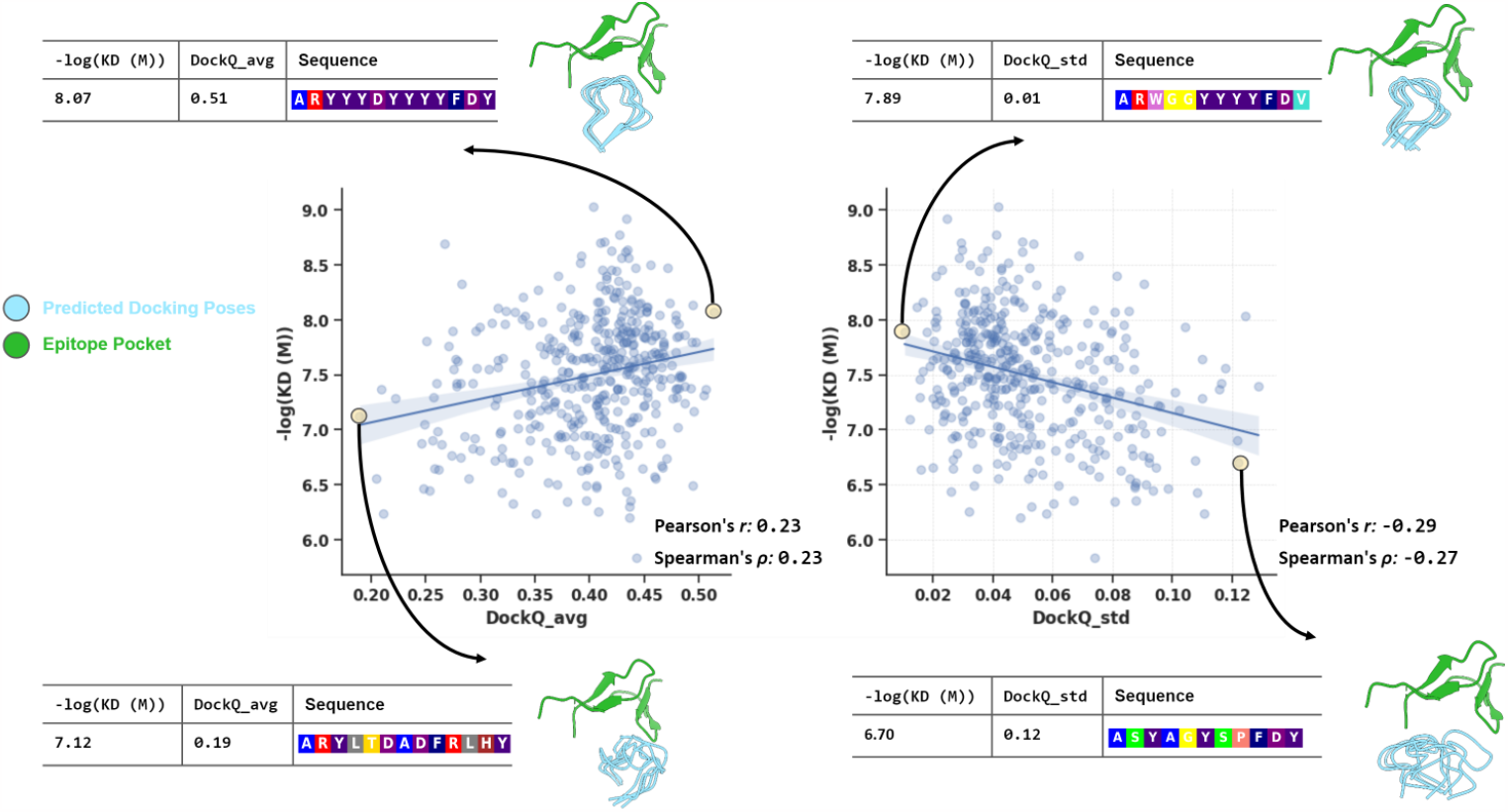
AbDock-Based Virtual Screening. Scatter plots depict the correlation between DockQ scores and antibody binding affinity (-log(KD(M))) for 422 CDRH3 variants targeting HER2. Positive correlation (left) suggests higher DockQ_avg values indicate stronger binding, while negative correlation (right) reveals lower DockQ_std values indicate higher prediction confidence. Highlighted cases showcase binding affinity scenarios.

**Fig. 6.**
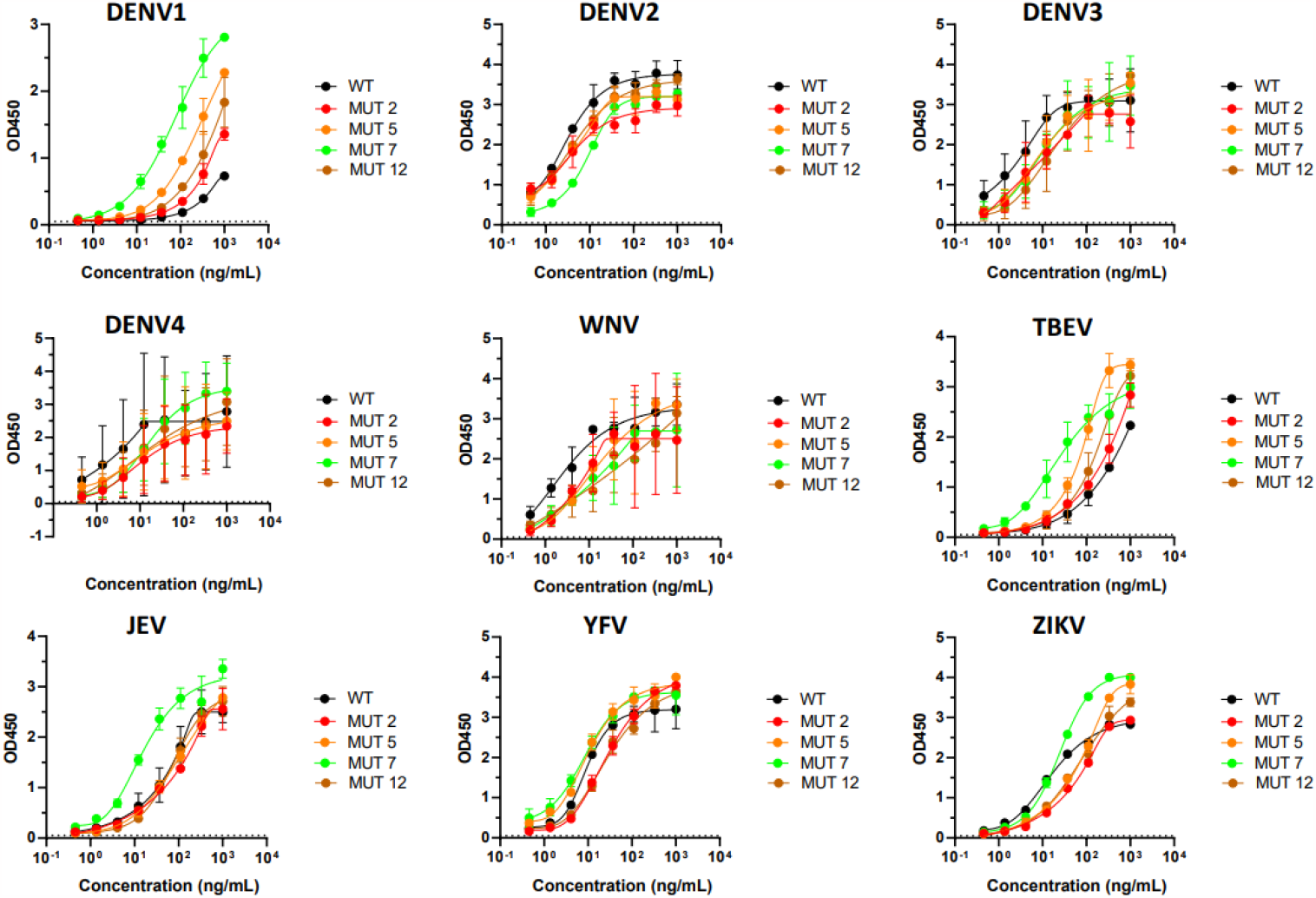
1G5.3 mAbs cross-reacts with full-length NS1 from DENV-1, DENV-2, DENV-3, DENV-4, WNV, TBEV, JEV, YFV, ZIKV. ELISA binding curves of purified flavivirus NS1 proteins coated on plates and detected with serial diluted purified 1G5.3 wild type (black), 1G5.3 mutant 2 (red), 1G5.3 mutant 5 (orange), 1G5.3 mutant 7 (green), 1G5.3 mutant 12 (brown) human monoclonal antibodies.

Through this process, we obtained a comprehensive in-silico score distribution for the designed antibody CDRH3 variants, allowing us to evaluate their potential binding affinities to the HER2 target. The utilization of AbDock in virtual screening enabled us to efficiently and reliably assess a large pool of antibody candidates, significantly reducing the time and resources required for experimental screening.

### 2.5 Optimizing Monoclonal Antibody 1G5.3 for Enhanced Binding Affinity and Broad-Spectrum Reactivity

To showcase the efficacy of our antibody optimization pipeline, we have applied the said pipeline to enhance the binding affinity of the Monoclonal antibody 1G5.3. This particular antibody, 1G5.3, presents a formidable challenge in terms of optimization due to its demonstrated exceptional affinity and broad cross-reactivity with the Flavivirus nonstructural protein 1 (NS1). To address this challenge and improve the affinity of the antibody, our approach involves the identification of four critical residues (118-122) within the CDRH3 that exhibit weak binding. These identified residues are then earmarked for redesign.

Subsequently, the optimization pipeline is deployed to facilitate the generation of 1,000 binding poses, alongside the redesigning of 1,000 sequence variants for these four designated residues. Next, virtual scoring is employed to evaluate the designed sequences. Based on the established criteria, which entails achieving a position within the top 50% of scores in DockQ_avg, as well as securing a position within the bottom 50% of scores in DockQ_std, a total of nineteen candidates are meticulously chosen for subsequent experimental validation. Among nineteen 1G5.3 mutant mAbs with the highest score screened using our model, we obtained four 1G5.3 mutants with high yield: mutant 2 (named MUT2 hereafter), MUT5, MUT7 and MUT12. To measure the cross-reactivity of these 1G5.3 mutants against NS1 proteins from different flaviviruses, we performed an enzyme-linked immunosorbent assay (ELISA) with purified mAbs and nine different flavivirus NS1 proteins. We tested the binding of the mAbs to four serotypes of dengue virus (DENV-1, DENV-2, DENV-3, DENV-4), as well as West Nile virus (WNV), tick-borne encephalitis virus (TBEV), Japanese encephalitis virus (JEV), yellow fever virus (YFV), and Zika virus (ZIKV). According to the results of the binding assays, some 1G5.3 mutants had improved specificity for different flaviviruses. Particularly, MUT2, MUT5, MUT7 and MUT12 exhibited higher binding activities than the wild-type 1G5.3 for DENV-1 and TBEV. Furthermore, binding for YFV and ZIKV were also increased in MUT5 and MUT7, and MUT7 had an additional specificity for DENV-4 and JEV. These results imply that the mutations of 1G5.3 antibody’s CDRs may affect how it recognizes the flavivirus NS1 proteins and confer a wider cross-reactivity. In conclusion, our pipeline successfully designed a broad-spectrum protective antibody 1G5.3 MUT7 monoclonal antibody which exhibited outstanding broad-spectrum binding to 6 out of 9 flaviviruses tested, indicating their potentials for universal flavivirus immunotherapy and vaccine design.

## 3 Conclusion

In this work, we present a diffusion model-based approach for antibody optimization with two key components: AbDesign and AbDock for antibody design and screening, respectively. Compared with previous algorithms, our models build up on the SE(3)-equivariant and -invariant neural network for modeling antibody-antigen complexes and generative diffusion models for sampling diverse candidates. By integrating the two techniques, our AbDesign and AbDock achieve state-of-the-art performance in independent test sets for design and docking tasks. In addition, the generative nature allows the models to design large-scale antibody candidates for further screening, aligning with the practical antibody optimization scenario. Furthermore, AbDock, specializing in paratope-epitope docking, offers a reliable measure of antibody quality by assessing generated docking samples, eliminating the need for costly specific binding affinity data. The in-silico antibody optimization framework is further experimentally validated through the optimization of monoclonal antibody 1G5.3. By designing and screening four residues on the CDRH3 of 1G5.3, we successfully obtained a variant that exhibited an exceptional and expansive binding capacity, effectively targeting six out of the nine tested flaviviruses.

While our pipeline has proven effective, it is important to acknowledge its inherent limitations. One notable limitation is the unsatisfactory scoring accuracy of our AbDock, which stands at Pearson’s correlation of 0.29. There is room for improvement in order to reduce errors in the screening stage. Additionally, our framework relies on the availability of antibody-antigen complex structures as a starting point for optimization. However, experimentally determining these complex structures can be a challenging task, particularly in cases where the binding between the antibody and antigen is weak, as is often encountered in optimization scenarios. Recent advancements in deep learning techniques have significantly improved the accuracy of protein structure prediction [26]. In situations where the native protein structure is unavailable, one potential solution is to utilize predicted complex structures [34]. However, these predicted complex structures often suffer from inaccuracies, particularly in the interface region, which, in turn, introduce errors to the optimization process. As a result, future research efforts should focus on optimizing our approach by considering scenarios where only the antigen pocket structure or sequence is available, thereby reducing the reliance on complete antibody-antigen complex structures. This would address the practical challenges associated with obtaining these complex structures and further enhance the applicability and robustness of our methodology.

## 4 Methods

In this work, we present the AbDesign for CDR design and AbDock for epitope-paratope docking as our proposed models. Although these models are designed for different tasks, they share the same diffusion formulation. In this section, we will begin by defining the problem setting for both the design and docking tasks. Next, we will introduce the diffusion process for modeling antibody structures. Lastly, we will describe the two neural networks utilized in our AbDesign and AbDock, respectively.

### 4.1 Problem Setting

Antibodies and antigens are protein entities composed of a sequence of amino acids denoted as {*R*_1_, …, *R*_*N*_ }, where *N* represents the number of amino acids in a protein. Each amino acid is characterized by its type *s*_*i*_ ∈ {*ACDEFGHIKLMNPQRSTV WY*}, *C*_*α*_ atom coordinate *x*_*i*_ ∈ ℝ^3^, and backbone orientation **O**_*i*_ ∈ *SO*(3). Here, *i* denotes the index of the amino acid within the protein, and we express it as *R*_*i*_ = {*s*_*i*_, *x*_*i*_, **O**_*i*_ }.

In the task of CDR sequence-structure co-design, given antibody framework and antigen structure, the model designs the CDRH3 (highlighted by a question mark in Fig. 1 B). We assume an antibody-antigen complex composed of N residues, encompassing the antibody framework, CDR, and antigen. More specifically, the first l residues correspond to the antibody framework, the middle m residues represent the CDR, and the remaining residues correspond to the antigen. The CDR to be generated consists of m amino acids, indexed from l+1 to l+m, denoted as ℛ = {(s_*j*_, **x**_*j*_, **O**_*j*_) | *j* = *l* + 1, …, *l* + *m*}. The task is to predict ℛ given the antibody framework and antigen residues *𝒞* = (*s*_*i*_, ***x***_*i*_, ***O***_*i*_) | *i* ∈ {1 … *l*}∪{*l* + *m*, …, *N*}}.

In the epitope-paratope docking task, we assume the knowledge of the antigen and the paratope sequence. The objective is to simultaneously fold the paratope sequence and dock it onto the antigen (Fig. 1 C). Similarly, the CDR structure to be generated is denoted as *𝒰* = {(**x**_*j*_, **O**_*j*_) | *j* = *l* + 1, …, *l* + *m*}. The task is to predict *𝒰* given the paratope sequence *𝒱* = {*s*_*i*_ | *i* ∈ {*l* + 1 … *l* + *m*}} and the antigen *𝒲* = {(*s*_*i*_, ***x***_*i*_, ***O***_*i*_) | *i* ∈ {*l* + *m*, …, *N* }}.

### 4.2 Diffusion Process

Diffusion models encompass a forward process wherein a sample is gradually perturbed by noise, while a neural network is trained to restore the original signal. Once the network is trained, the sample can be generated by progressively removing noise from an initial noisy state. This section introduces the diffusion processes (forward and backward) employed in protein structure design, specifically focusing on the *C*_*α*_ coordinates and backbone orientations. The sequence design is conducted by directly predicting the amino acid type of each residues instead of using diffusion models, as introduced in Section 4.3.1.

#### 4.2.1 Diffusion for *C*_*α*_ Coordinates

We utilize the original diffusion models [22] to establish the diffusion process on *C*_*α*_ coordinates. The forward process involves the interpolation between the *C*_*α*_ coordinates and the standard Gaussian distribution. It can be formally defined as follows:

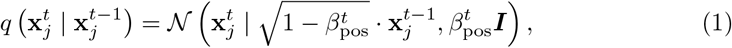

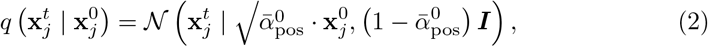

where 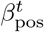 represents a predefined noise schedule that regulates the noise ratio. It gradually increases from 0 to 1 as the time step progresses from 0 to T. Additionally 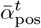, is computed as the product of 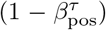 over *τ* from 1 to t. Here, the index *j* denotes the diffused residues.

The backward process, based on the reparameterization trick proposed by [22], can be defined as follows:

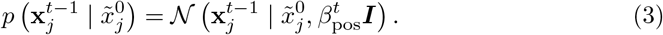

Here, 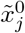 represents the predicted *C*_*α*_ coordinates for residue *j* at time step 0, obtained through the neural network *f*_*θ*_:

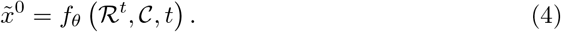

Here *θ* denotes the learnable parameter of the neural network *f* (.). ℛ^*t*^ corresponds to the diffused residues at time *t*, where *t* ∼ *U* [1, *T*].𝒞 denotes the contextual interface. The objective function for training the generative process involves the expected mean squared error (MSE) between the predicted *C*_*α*_ coordinates 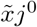and the ground truth *Cα* coordinates 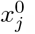. It can be expressed as:

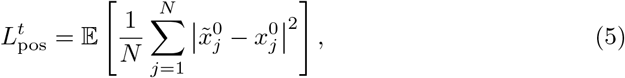

In the above equation, *N* represents the total number of diffused residues. The objective is to minimize this expected MSE during training.

#### 4.2.2 nSO(3) Diffusion for Backbone Orientations

The residue backbone orientation can be mathematically represented as rotation in 3D Euclidean space, which is elements of SO(3) group. We formulate the forward process according to [10, 35]:

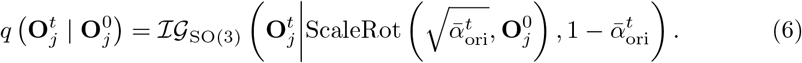

The distribution ℐ 𝒢SO (3) represents an isotropic Gaussian distribution on the SO(3) group, which is parameterized by a mean rotation and a scalar variance [35, 36]. The ScaleRot operation modifies the rotation matrix by scaling its rotation angle while keeping the rotation axis fixed [37]. We define 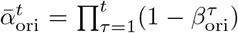 where 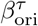 is a pre-specified noise schedule.

The backward process is defined as:

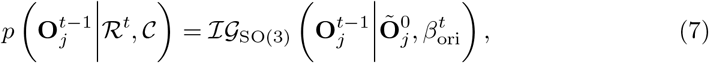

where 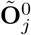 is the orientation of residue i at time step 0, obtained using the neural network *g*_*θ*_(*·*):

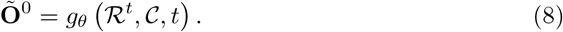

The training objective aims to align the predicted orientation with the ground truth by minimizing the expected discrepancy, which is measured using the inner product between the real and predicted orientation matrices. The corresponding loss function is defined as follows:

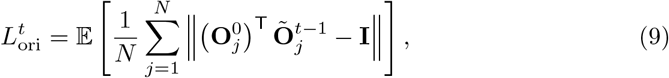

where **I** is the 3 *×* 3 identity matrix.

#### 4.2.3 Training Algorithms

To train the networks 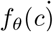 and 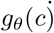, we follow the procedure below. We uniformly sample time step *t* from *t ∼ U* [1, *T*] and perform the forward process by adding noise to the native *C*_*α*_ coordinates and orientation. These perturbed samples are then fed into the networks to obtain the predicted samples 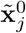 and 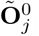. Next, we calculate the losses between native and predicted samples and update the parameters *θ* using gradient descent with the Adam optimizer. In addition to the aforementioned losses, we incorporate a distance loss to optimize the prediction of *Cα* coordinates, defined as follows:

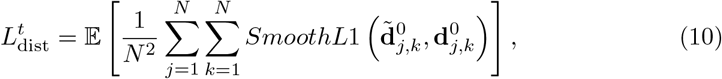

The term 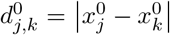 the Euclidean distance between *C*_*α*_ coordinates 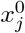 and 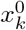. The *SmoothL*1 function applies a squared term when the absolute elementwise error is less than 1, and an *L*1 term otherwise. The diffusion loss is computed as the sum of three terms:

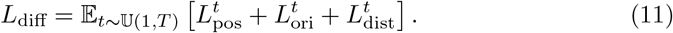

In sequence design, we utilize a cross-entropy loss function to optimize the prediction of amino acid types, denoted as *L*_seq_. This loss term contributes to the overall loss for the sequence structure co-design task, which can be expressed as:

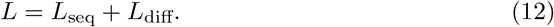

#### 4.2.4 Sampling Algorithms

To generate the CDR structures, we follow the procedure outlined below. Firstly, we sample the 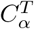 coordinates and orientation **O**^*T*^ from a prior distribution: 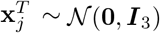 and 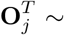Uniform(*SO*(3)). Subsequently, we iteratively run the backward processes until *t* = 1 to obtain obtain the 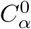and **O**^0^.

To construct the complete atomistic 3D structure, we determine the coordinates of the N, *C*_*α*_, C, O, and side-chain *C*_*β*_ atoms, excluding glycine which lacks a *C*_*β*_ atom. These atoms are positioned according to their ideal local coordinates relative to the *C*_*α*_ position and orientation of each amino acid, based on the reconstruction of the five key atoms [26]. The remaining side-chain atoms are generated using the side-chain packing function implemented in Rosetta [38]. Finally, we refine the full atom structure using the AMBER99 force field [39] within the OpenMM framework [32].

### 4.3 Neural Network Architectures

To address the specific demands of the design and docking tasks, we leverage distinct network architectures (Fig. 1 D). For the design task, we utilize the Multi-Channel Equivariant Graph Neural Network (MC-EGNN) implemented in AbDesign. Conversely, in the docking task, we employ Invariant Point Attention (IPA) as part of the AbDock framework.

#### 4.3.1 Multi-Channel Equivariant Graph Neural Network

We represent the binding interface between antibodies and antigens as a locally connected graph and employ the Multi-Channel Equivariant Graph Neural Network (MC-EGNN) [23, 25] (Fig. 1 D left) to capture the geometry and topology of this interface. The node features in the graph encode information such as amino acid types, torsional angles, and 3D coordinates of all heavy atoms. The graph’s edges consist of internal and external connections. Internal edges are constructed between residues within the same chain if their *C*_*α*_ distance is below 8Å, while external edges are established between residues in different chains if their *C*_*α*_ distance is below 12Å.

MC-EGNN leverages the E(3)-equivariant message passing mechanism to update the internal and external graphs separately. Through multiple layers of message passing, the node representations and coordinates are transformed into predictions for amino acid types, 3D positions, and orientations of the complementary determining regions (CDRs). MC-EGNN comprises an internal context encoder to update the internal graph and an external attentive encoder to update the external graph. The internal context encoder employs multi-layer perceptrons (MLPs) to generate messages between nodes based on their node embeddings and the relative distances of backbone atoms. The node embeddings and coordinates are further updated by passing the messages through two MLPs. The external attentive encoder utilizes a graph attention mechanism to describe the correlation between distant residues in an E(3)-equivariant manner. This module generates query, key, and value vectors, as well as attention weights, from the node embeddings. The node embeddings and coordinates are then updated by applying the attention weights to the values. After 6 layers of alternating between the internal and external encoder, we feed the node features to an internal context encoder to predict the amino acid types and coordinates.

#### 4.3.2 Invariant Point Attention

We employ the Invariant Point Attention (IPA) [26] ((Fig. 1 E right) to capture long-range interactions in paratope-epitope complexes and predict paratope binding poses. Initially, we utilize Multiple Layer Perceptrons (MLPs) to generate single and pair representations from the given paratope sequence *𝒱* and antigen *𝒲*. The single representation, denoted as *e*_*j*_, encodes information about amino acid types, torsional angles, and 3D coordinates of all heavy atoms. Here, *e*_*i*_ represents the embedding for the *i*-th amino acid, and *N* denotes the total number of amino acids. On the other hand, the pair representation, denoted as *z*_*i,j*_, encodes pairwise interactions between two amino acids. In particular, *z*_*i,j*_ encodes Euclidean distances and dihedral angles between amino acid *i* and *j*.

To update the hidden representation, we employ a 6-layer IPA. the IPA combines the pair representation, the single representation, and the geometric representation to update the single representation. The IPA operates in 3D space, where each residue produces query points, key points, and value points in its local frame. These points are projected into the global frame using the backbone frame of the residue in which they interact with each other. The resulting points are then projected back into the local frame. The affinity computation in the 3D space uses squared distances, and the coordinate transformations ensure the invariance of this module with respect to the global frame. The IPA is an equivariant operation on the residue gas, and it imposes a strong spatial/locality bias on the attention, which is well-suited to the iterative refinement of the protein structure. This integration of multiple representations allows the IPA to effectively update the single representation, incorporating both local and global structural information. Finally, we feed the updated single representation into two separate MLPs to denoise the 3D positions and orientations of the paratope, respectively.

### 4.4 Antibody Optimization pipeline

In this section, we present the antibody optimization pipeline utilizing the proposed AbDesign and AbDock methods. Given an antibody-antigen complex, the goal of antibody optimization is to enhance the antibody binding affinity by redesigning its sequence. Traditionally, a two-step pipeline involving design and screening might seem more straightforward. However, we have observed that this approach often lacks sequence diversity. To address this issue and improve sequence diversity, we propose a three-step pipeline (Fig. 1 E) to generate a broader range of sequences for further screening. The process involves three main steps: binding pose generation, sequence design, and screening. First, we employ AbDock to generate different binding poses by inputting the native CDR sequences and sampling multiple docking poses. Next, we use AbDesign (seq. design version) to redesign the CDR sequences based on the given complex structures and the generated poses. Finally, the designed sequences undergo in-silico screening, where AbDock is employed again to score and rank them based on generated metrics: DockQ_avg and DockQ_std.

### 4.5 IgG binding ELISA

For ELISA binding assays of 1G5.3 mutants, the antigen panel included DENV1-NS1, DENV2-NS1, DENV3-NS1, DENV4-NS1, WNV-NS1, TBEV-NS1, JEV-NS1, YFV-NS1, ZIKV-NS1 from human cell Expi293F™ Cells (ThermoFisher, Catalog no. A14527). For binding ELISA, 96-well ELISA plates were coated with 1 μg/mL of antigens in 1X PBS (Macgene, Catalog no. CC008) overnight at 4°C. Plates were washed with PBS + 0.01% Tween 20 and blocked with assay diluent (150mM NaCl, 50mM Tris-HCl, 1mM EDTA, 3.3% FBS, 2% BSA, 0.07% Tween-20) at 37 degrees for 2 hours. Purified mAb samples in 3-fold serial dilutions in assay diluent starting at 1 μg/mL and incubated at 37 degrees for 2 hours, followed by washing with PBS + 0.01% Tween 20 (Aladin, Catalog no. 9005-64-5). Anti-Human IgG (H+L) HRP Conjugate (Promega, catalog no. W4031) was diluted to 1:5,000 and incubated at 37 degrees for 1 hour. These plates were washed five times with PBS + 0.01% Tween 20 and developed with TMB detection Kit (CW Bio Catlog no. CW0050S). The reaction was stopped with ELISA stop solution (Solarbio Catalog no. C1058), and optical density at 450 nm (OD450) was determined using a microplate reader (BioTek Synergy H1).

## 5 Data Availability

The datasets utilized in this project are publicly available. The antibody-antigen structural data used for training was sourced from the SAbDab database https://opig.stats.ox.ac.uk/webapps/newsabdab/sabdab/. The binding affinity data used for evaluating the scoring performance of AbDock is available at https://github.com/AbSciBio/unlocking-de-novo-antibody-design.

## 6 Code Availability

The code for reproducing the training and testing results is made available at https://github.com/pengzhangzhi/ab_opt. A website animating the generation trajectories is available at https://pengzhangzhi.github.io/ab_opt_homepage/.

## Footnotes

1 https://github.com/AbSciBio/unlocking-de-novo-antibody-design

2 https://www.rcsb.org/structure/1N8Z

